# Benzoate and Salicylate Tolerant Strains Lose Antibiotic Resistance during Laboratory Evolution of *Escherichia coli* K-12

**DOI:** 10.1101/063271

**Authors:** Kaitlin E. Creamer, Frederick S. Ditmars, Preston J. Basting, Karina S. Kunka, Issam N. Hamdallah, Sean P. Bush, Zachary Scott, Amanda He, Stephanie R. Penix, Alexandra S. Gonzales, Elizabeth K. Eder, Dominic Camperchioli, Adama Berndt, Michelle W. Clark, Kerry A. Rouhier, Joan L. Slonczewski

**Author notes:** Corresponding Author: Joan L. Slonczewski Robert A. Oden, Jr. Professor of Biology Higley Hall, 202 N. College Road Kenyon College Gambier, OH 43022 http://biology.kenyon.edu/slonc/slonc.htm. These authors contributed equally to the work (Kaitlin E. Creamer, Frederick S. Ditmars, Preston J. Basting).

## Abstract

*Escherichia coli* K-12 W3110 grows in the presence of membrane-permeant organic acids that can depress cytoplasmic pH and accumulate in the cytoplasm. We conducted experimental evolution by daily diluting cultures in increasing concentrations of benzoic acid (up to 20 mM) buffered at external pH 6.5, a pH at which permeant acids concentrate in the cytoplasm. By 2,000 generations, clones isolated from evolving populations showed increasing tolerance to benzoate but were sensitive to chloramphenicol and tetracycline. Sixteen clones grew to stationary phase in 20 mM benzoate, whereas the ancestral strain W3110 peaked and declined. Similar growth occurred in 10 mM salicylate. Benzoate-evolved strains grew like W3110 in the absence of benzoate; in media buffered at pH 4.8, pH 7.0, or pH 9.0; or in 20 mM acetate or sorbate at pH 6.5. Genomes of 16 strains revealed over 100 mutations including SNPs, large deletions, and insertion knockouts. Most strains acquired deletions in the benzoate-induced multiple antibiotic resistance (Mar) regulon or in associated regulators such as *rob* and *cpxA*, as well as MDR efflux pumps *emrA*, *emrY*, and *mdtA*. Strains also lost or down-regulated the Gad acid fitness regulon. In 5 mM benzoate, or in 2 mM salicylate (2-hydroxybenzoate), most strains showed increased sensitivity to the antibiotics chloramphenicol and tetracycline; some strains were more sensitive than a *marA* knockout. Thus, our benzoate-evolved strains may reveal additional unknown drug resistance components. Benzoate or salicylate selection pressure may cause general loss of MDR genes and regulators.

**IMPORTANCE:** Benzoate is a common food preservative, and salicylate is the primary active metabolite of aspirin. In the gut microbiome, genetic adaptation to salicylate may involve loss or downregulation of inducible multidrug resistance systems. This discovery implies that aspirin therapy may modulate the human gut microbiome to favor salicylate tolerance at the expense of drug resistance. Similar aspirin-associated loss of drug resistance might occur in bacterial pathogens found in arterial plaques.

## INTRODUCTION

Pathogenic and commensal enteric bacteria maintain cytoplasmic pH homeostasis in the face of extreme external acid (pH 2–4 in the stomach) and the high concentrations of membrane-permeant organic acids in the colon (70–140 mM) (1–6). Many studies have focused on the response and recovery of *E. coli* to external pH stress (1–5), but relatively few studies have focused on the genetic response to membrane-permeant organic acids (permeant acids) (7–9) despite the importance of permeant acids as food preservatives (10). Permeant acids depress cytoplasmic pH, while their anion accumulates in the cytoplasm (5, 7, 11). Some permeant acids also cross the membrane in the unprotonated form, and decrease the proton motive force (PMF); an example is benzoic acid (benzoate), a food preservative found in soft drinks and acidic foods (12). The related molecule salicylic acid (salicylate) is a plant defense regulator (13, 14) as well as the primary active metabolite of acetylsalicylate (aspirin) (15–17). Salicylates enter the human diet from fruits and vegetables, leading to circulating plasma levels as high as 0.1-0.2 μM (18). Furthermore, aspirin therapy for cardio protection and other metabolic conditions (19, 20) may generate plasma levels of 0.2-0.5 mM salicylate (16, 21, 22). Yet despite the important metabolic effects of aspirin, salicylate and benzoate on plants and animals, there is surprisingly little research on the effects of benzoate and salicylate on the host microbiomes. In one study, aspirin inhibits the growth of *Helicobacter pylori* and enhances the pathogen’s sensitivity to antibiotics (23).

In *E. coli*, aromatic permeant acids such as salicylate and benzoate induce a large number of low-level multidrug efflux systems, governed by the Mar operon (*marRAB*) as well as additional unidentified mechanisms (24). Benzoate and salicylate upregulate numerous genes of commensals and pathogens (25–28) including *acrAB*, *tolC*, and transport complexes that expel drugs across both the cytoplasmic and outer membrane. Mar-family systems are widespread in bacterial genomes (29). Thus, in natural environments, aromatic acids may serve bacteria as early warning signals for the presence of antibiotic-producing competitors.

Mar-family regulons commonly involve extensive upregulation of many genes by a small number of regulators. The *E. coli* regulator MarR represses expression of *marRAB*; repression is relieved when MarR binds salicylate (30) or one of several less potent inducers such as benzoate or 2,4-dinitrophenol. The upregulated MarA is an AraC-type global regulator that differentially regulates approximately 60 genes (27, 31). Another AraC-type regulator, Rob, activates *marRAB* (26, 32). MarA downregulates the acid-inducible Gad acid fitness island (33). Gad includes glutamate decarboxylase (*gadA*) for extreme-acid survival (32–34), as well as periplasmic chaperones *hdeA* and *hdeB* (35), and MDR loci *mdtE*, *mdtF* (36). Besides Mar, short-term benzoate exposure up-regulates biofilm-associated genes (*ymgABC*, *yhcN*), the fimbrial phase-variation regulator (*fimB*), and the cadmium stress protein *yodA* (37).

Thus, aromatic acid-inducible drug resistance incurs high energy costs associated with expression of so many genes, as well as the energy consumption by efflux pumps (38). Given the high energy cost, bacteria face a tradeoff between inducible drug resistance and the toxicity of the drugs (39). One would expect a high selective pressure for regulator alleles that shift expression based on environmental factors. In fact, plate-based selection screens using *lac* fusions readily pick up mutations in *marR* and in MarR-regulated genes (8). Selective growth under antibiotic pressure leads to upregulation of *marRAB* (40).

A powerful tool for dissecting long-term response to environmental stresses is experimental evolution (41, 42). Experimental evolution procedures with *E. coli* have included the adaptation to high temperatures (43), freeze-thaw cycles (44), high ethanol concentrations (45), and acid (46–48). We developed a microplate dilution cycle in order to generate evolving populations buffered at low pH (49). An advantage of our microplate dilution cycle is that we propagate a number of populations directly in the microplate, eliminating the intermediate stage of culture in flasks or tubes.

For the present study, we conducted experimental evolution of *E. coli* K-12 W3110 in microplate well populations containing media buffered at pH 6.5 and supplemented with increasing concentrations of benzoate (from 5 mM initially to 20 mM at 2,000 generations). We sequenced genomes of selected isolates, then identified genetic variants using the *breseq* pipeline (48–50). The *breseq* pipeline assembles a reference-based alignment to predict mutations compared to a previously sequenced genome (NCBI GenBank accession number NC_007779.1, *E. coli* K-12 W3110). Newer versions of *breseq* now predict structural variations including large deletions, mobile element insertions, and gene duplications—all of which account for much of the genetic diversity in evolved clones (50–53).

Our analysis unexpectedly shows that genetic adaptation to benzoate is associated with loss or down-regulation of benzoate- and salicylate-inducible genes, including those that encode multidrug resistance systems. The results have implications for evolution of the gut microbiome during aspirin therapy. More broadly, our results suggest a way to amplify the fitness costs of antibiotic resistance and possibly reverse antibiotic resistance in a microbiome (54).

## MATERIALS AND METHODS

### Bacterial strains and media

*Escherichia coli* K-12 W3110 (55) was the ancestral strain of all benzoate-adapted populations. Additional strains derived from *E. coli* K-12 W3110 were isolated during the course of the evolution experiment (**Table 1**). Alleles with *kanR* insertions were obtained from the Keio Collection (56) distributed by the Coli Genetic Stock Center (CGSC). A *marR*::*kanR* strain was provided by Frederick R. Blattner, University of Wisconsin-Madison.

**Table 1.**
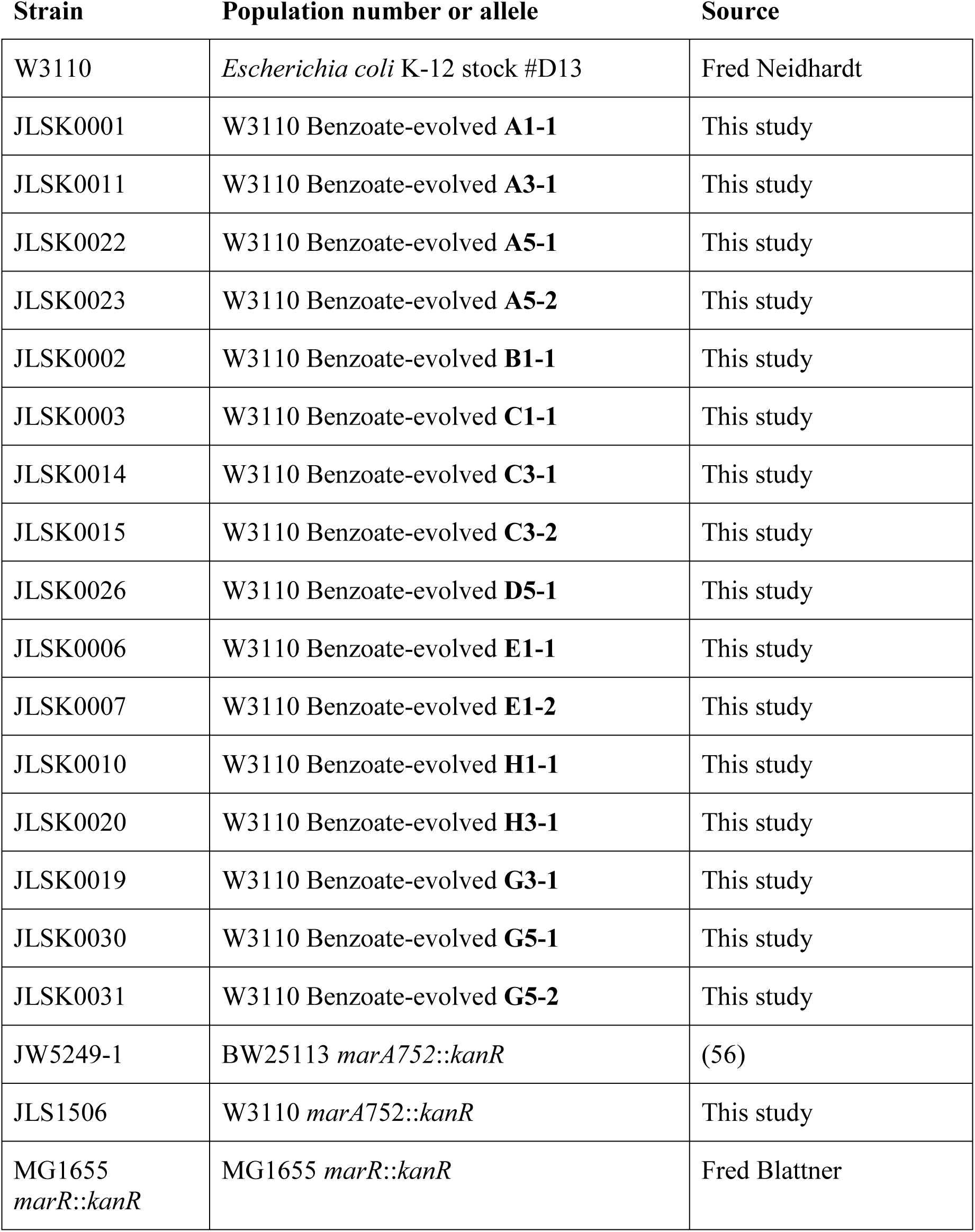
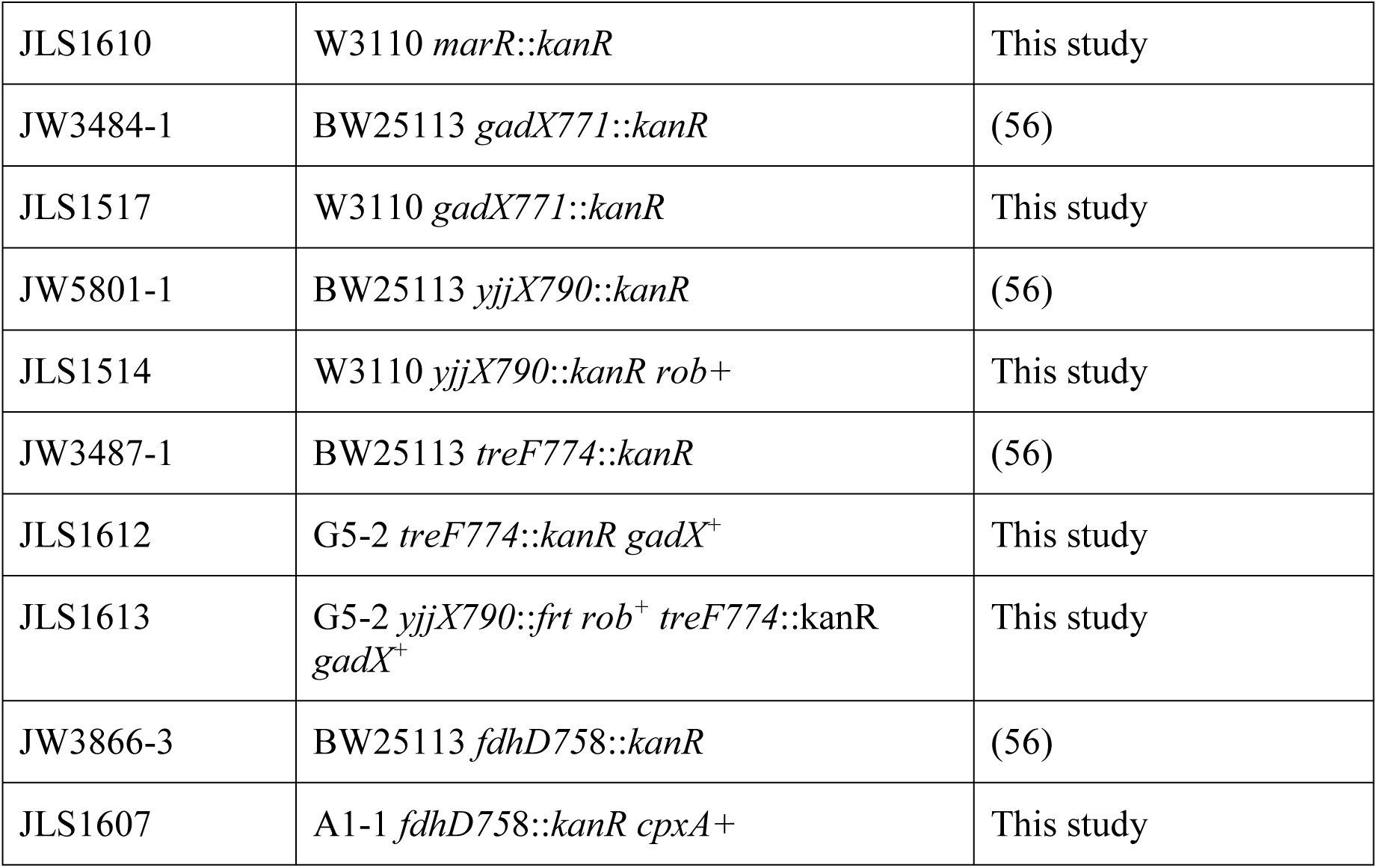
**Bacterial strains used in this study.**

Bacteria were cultured in LBK (10 g/L tryptone, 5 g/L yeast extract, 7.45 g/L KCl) (57). Culture media were buffered with either 100mM piperazine-N,N’-bis(ethanesulfonic acid) (PIPES; pKa= 6.8), 100 mM, 2-(N-morpholino)ethanesulfonic acid (MES; pKa= 5.96), 100 mM 3-morpholinopropane-1-sulfonic acid (MOPS; pKa= 7.20), 100 mM homopiperazine-N,N’-bis-2-(ethanesulfonic acid) (HOMOPIPES; pKa = 4.55, 8.12), or 150 mM N-Tris(hydroxymethyl)methyl-3-aminopropanesulfonic acid (TAPS; pKa= 8.4). The pH of the medium was adjusted as necessary with either 5 M HCl or 5 M KOH. Potassium benzoate (referred to as benzoate), sodium salicylate (salicylate), potassium acetate, potassium sorbate, chloramphenicol, or tetracycline was added before filter sterilization for LBK media requiring various concentrations of acids or antibiotics. Temperature of incubation was 37°C unless noted otherwise.

### Experimental Evolution

Experimental evolution was conducted according to the procedure of our low-pH laboratory evolution experiment (49) with modifications. Briefly, 24 cultures derived from the same ancestral strain (W3110, freezer stock D13) were cultured continuously in increasing concentrations of benzoate for 2,000 generations (**Figure S1**). An overnight culture of ancestral *Escherichia coli* K-12 W3110 was diluted 1:100 in LBK, pH 6.5 100 mM PIPES, 5 mM potassium benzoate. Growth was recorded over 22 h in a SpectraMax Plus384 MicroPlate reader (Molecular Devices). Every 15 minutes, the microplate was shaken for 3 seconds and the OD_450_ of each culture was recorded. The cultures were re-diluted 1:100 into fresh benzoate growth medium at the end of the daily cycle. 100 μl glycerol (50% glycerol, 100 mM MES pH 6.5) was added to each well, after which the microplate was frozen at −80°F (49). The number of generations of the exposed cells was calculated based on the 1:100 daily dilution, resulting in a 100-fold daily growth to achieve approximately 6.6 generations of binary fission (58). In the course of the 22-hour cycle, all bacterial populations attained stationary phase densities. Plating representative cultures showed that during the dilution cycle cell numbers increased from approximately 5×10^6^ cells per ml to 5×10^8^ cells per ml.

Over the course of the experiment, the benzoate concentration was increased in steps when all populations of the plate had attained approximately100-fold stationary growth over multiple dilution cycles (see **Figure S1**). After 60 generations, the benzoate was increased to 6 mM; 10 mM after 90 generations; 12 mM after 540 generations; 15 mM after 1,020 generations; 18 mM after 1,210 generations; 20 mM after 1,580 generations, to the conclusion of the experiment with a cumulative 3,000 generations of growth. If the strains had to be restarted from a frozen microplate, the frozen cultures were thawed and diluted 1:50 into fresh potassium benzoate growth media.

After 2,000 generations, microplates were taken from the freezer and samples from specific wells were spread on LBK agar plates. Selected clones from each chosen well were streaked three times and stored as freezer stocks. Clones were cultured in media at pH 6.5, with 5 mM benzoate; all clones showed increased growth compared to ancestral strain W3110 (data not shown). For genome sequencing, eight clones were chosen in pairs from each of four populations (clones A5-1, A5-2, C3-1, C3-2, E1-1, E1-2, G5-1, G5-2). A total number of 24 clones (one from each population) were tested for sensitivity to chloramphenicol (8 μg/ml) in 5 mM benzoate medium, pH 7.0. From these, eight additional chloramphenicol-sensitive clones were selected for genomes sequencing (A1-1, A3-1, B1-1, C1-1, D5-1, G3-1, H1-1, H3-1). All sixteen strains are identified in **Table 1**; and their genomic mutations (compared to ancestral strain W3110) are presented in **Table S1**. Mutations in six selected strains are presented in **Table 2**. These strains are color-coded throughout our figures.

**Table 2.**
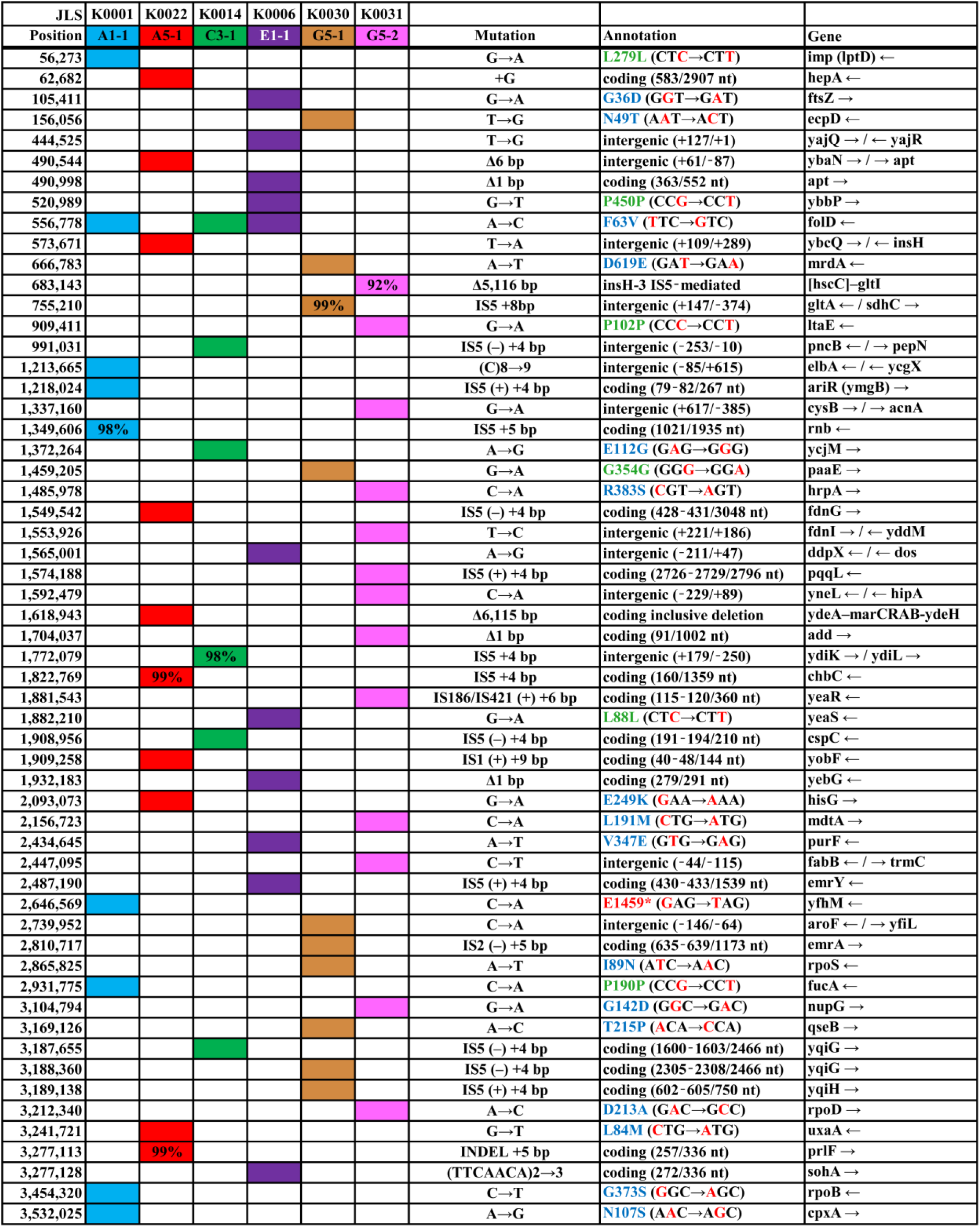
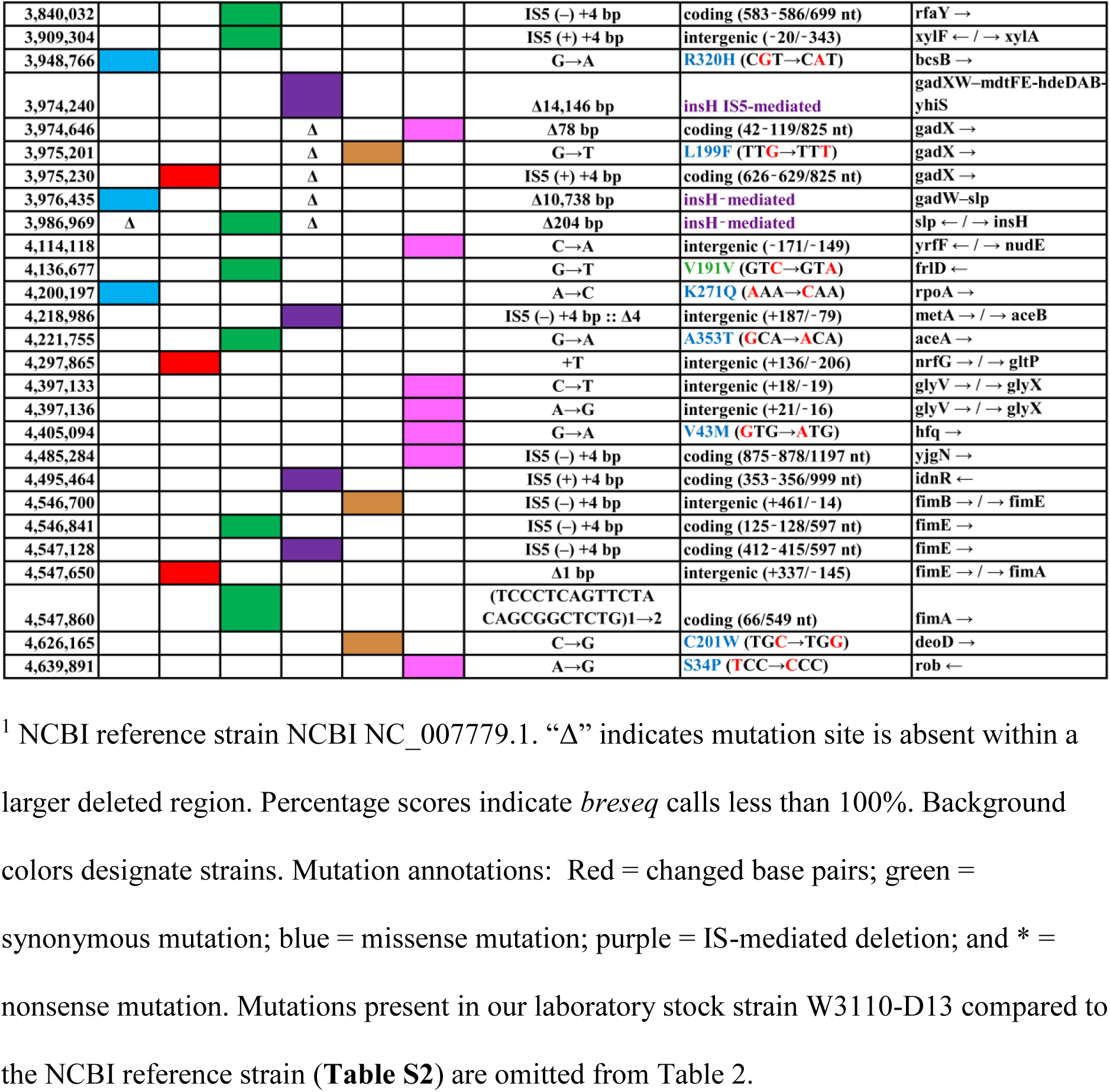
**Mutations in representative benzoate-evolved genomes compared to the genome of *E. coli* W3110**.

### Growth assays

Growth curves were measured in the microplate reader at 37°C for 22 hours under various conditions of organic acids, pH values, and antibiotics. Strains were cultured overnight in LBK pH 5.5 buffered with 100 mM MES; LBK pH 6.5 buffered with 100 mM PIPES; LBK pH 7.0 buffered with 100 mM MOPS; or LBK pH 8.5 buffered with 150 mM TAPS. Supplements included benzoate, salicylate, acetate, or sorbate, as stated in figures. Overnight cultures were diluted 1:100 (1:200 for the antibiotic growth assays) into the exposure media which included LBK pH 6.5 buffered with 100 mM PIPES; LBK pH 4.8, 100 mM HOMOPIPES; LBK pH 7.0, 100 mM MOPS; LBK pH 9.0, 150 mM TAPS; or LBK pH 7.0, 100 mM MOPS. Every 15 minutes, the plate was shaken for 3 seconds and an OD_600_ measurement was recorded. The growth rate *k* of each culture was calculated over the period of 1–3 h, approximately the log phase of growth (34). The cell density *E* of each culture was measured at 16 h unless stated otherwise.

### Genomic DNA extraction and sequencing

Genomic DNA from benzoate-evolved clones and from the ancestral wild type strain W3110 (freezer stock D13) was extracted using the DNeasy DNA extraction kit (Qiagen) and the MasterPure Complete DNA and RNA Purification Kit (Epicentre). The DNA purity was confirmed by measuring the 260nm/280nm and 260nm/230nm absorbance ratios using a NanoDrop 2000 spectrophotometer (Thermo Fisher Scientific) and the concentration of the DNA was measured using both the NanoDrop 2000 spectrophotometer (Thermo Fisher Scientific) and a Qubit 3.0 Fluorometer (Thermo Fisher Scientific), according to manufacturer instructions.

The genomic DNA was sequenced by Michigan State University Research Technology Support Facility Genomics Core. For Illumina MiSeq sequencing, libraries were prepared using the Illumina TruSeq Nano DNA Library Preparation Kit. After library validation and quantitation, they were pooled and loaded on an Illumina MiSeq flow cell. Sequencing was done in a 2×250 bp paired-end format using an Illumina 500 cycle V2 reagent cartridge. Base calling was performed by Illumina Real Time Analysis (RTA) v1.18.54 and output of RTA was demultiplexed and converted to FastQ format with Illumina Bcl2fastq v1.8.4.

### Nucleotide sequence accession number

Sequence data have been deposited in the NCBI Sequence Read Archive (SRA) under accession number SRP074501.

### Sequence assembly and analysis using *breseq* computational pipeline

The computational pipeline *breseq* version 0.27.1 was used to assemble and annotate the resulting reads of the evolved strains (50–52). The current *breseq* version detects IS element insertions and IS-mediated deletions, as well as SNPs and other mutations (53). The reads were mapped to the *E. coli* K-12 W3110 reference sequence (NCBI GenBank accession number NC_007779.1) (59). Mutations were predicted by *breseq* by comparing the sequences of the evolved isolates to that of the ancestral strain W3110, lab stock D13 (52).

In order to visualize the assembly and annotations of our evolved isolate sequences mapped to the reference *E. coli* K-12 W3110 genome, we used Integrative Genomics Viewer (IGV) from the Broad Institute at MIT (60). Sequence identity of clones was confirmed by PCR amplification of selected mutations.

### P1 phage transduction and strain construction

P1 phage transduction was conducted by standard procedures to replace a linked mutation with the ancestral non-mutated allele, as well to construct knockout strains (49). Strains with *kanR* insertions (56) were introduced into the evolved strain of choice or the ancestral strain W3110. Constructs were confirmed by PCR amplification and Sanger sequencing of key alleles of donor and recipient.

### MIC assays

For assays of minimum inhibitory concentration (MIC) of antibiotics (chloramphenicol or tetracycline) the strains were cultured in a microplate for 22 h in LBK 100 mM MOPS pH 7.0, 2 mM salicylate. The medium contained a range of antibiotic concentration (μg/ml): 0, 1, 2, 4, 6, 8, 12, 16, 24. A positive result for growth was defined as measurement of OD_600_ ≥ 0.05. Each MIC was reported as the median value of 8 replicates. For each antibiotic, three sets of 8 replicates were performed; that is, a total of 24 replicates per strain.

### GABA assays

The procedure for measuring GABA production via glutamate decarboxylase was modified from that of Ref. (61). Strains were cultured overnight in LB medium (10 g/L tryptone, 5 g/L yeast extract, 100 mM NaCl) buffered with 100 mM MES pH 5.5. 10 mM glutamine was included, which the bacteria convert to glutamate, the substrate of glutamate decarboxylase (62). For anaerobic culture, closed 9-ml screwcap tubes nearly full of medium were incubated for 18 hours at 37°C. The pH of each sample was lowered with HCl to pH 2.0 for extreme-acid stress (63) and incubated 2 h with rotation. Cell density (OD_600_) was measured in microplate wells, in a SpectraMax plate reader. 1 ml of each culture was pelleted in a microfuge. The supernatant was filtered and prepared for GCMS by EZ:faast amino acid derivatization (64). GABA concentration was calculated using a standard solution prepared at a concentration of 200 nm/ml. GABA and other compounds from the culture fluid were identified using NIST library analysis. GABA concentrations were normalized to OD_600_ values of the overnight cultures before assay.

## RESULTS

### Experimental evolution of benzoate-tolerant strains

We conducted experimental evolution of *Escherichia coli* K-12 W3110 exposed to increasing concentrations of benzoic acid, as described under Methods (**Figure S1**). Over the course of the experiment, bacteria showed progressive increase of tolerance to benzoate tolerance, as measured by endpoint culture density. Clones were sampled from microplate populations frozen at intervals over the course of zero to 2,900 generations (**Fig. 1**). The clones were cultured in microplate wells in media containing 20 mM benzoate at pH 6.5, and the cell density was measured at 16 h. Over 1,400–2,900 generations the growth endpoints of evolved clones increased significantly compared to that of the ancestor (**Fig. 1A**). A similar increase was observed for cell density during growth with 10 mM salicylate (**Fig. 1B**). Thus overall, tolerance to benzoate and salicylate increased over generations of exposure.

**FIG 1.**
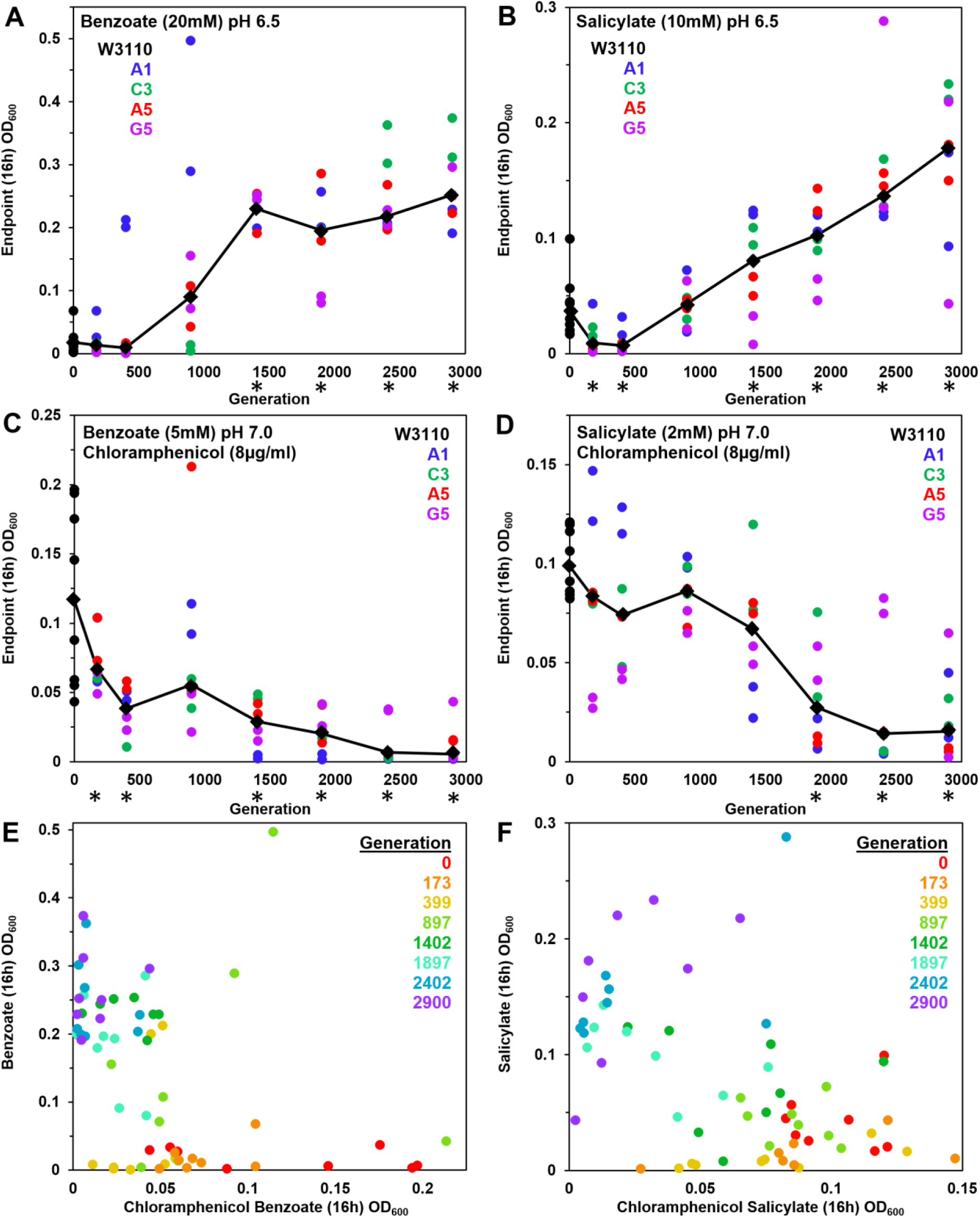
Growth measured after generations of repeated dilution and culture in 5–20 mM benzoate. Benzoate-evolved strains were isolated from frozen microplates, selecting 2 different clones from each of 4 populations. Clones from each plate generation were cultured at 37°C in a column of microplate wells; cell density “E” values (OD_600_) were obtained at 16 h. For panels A-D, the colored dots represent lineages of evolved clones collected from distinct well populations (A1, C3, A5, G5) after different numbers of generations as indicated on the X axis. Diamonds indicate median cell density for each generation tested. Bracket indicates generations for which the 16-h cell density differed significantly from that of the ancestral strain W3110, in 2 out of 3 trials of the entire microplate experiment. For each microplate trial, the Friedman test was performed with post-hoc Conover pairwise comparisons and Holm-Bonferroni adjusted p-values. LBK media contained: **A.** 100 mM PIPES pH 6.5 with 20 mM benzoate (diluted 1:200 from overnight cultures with 5mM benzoate). **B.** 100 mM PIPES pH 6.5 with 10 mM salicylate (diluted 1:200 from overnight cultures in 2 mM salicylate). **C.** 100 mM MOPS pH 7.0 with 5 mM benzoate, 8 μg/ml chloramphenicol (diluted 1:200 from overnight cultures lacking chloramphenicol). **D.** 100 mM MOPS pH 7.0 with 2 mM salicylate, 8 μg/ml chloramphenicol (diluted 1:200 from overnight cultures lacking chloramphenicol). **E.** Plot with linear regression of 16-h cell-density values for 20 mM benzoate and for 5 mM benzoate, 8 μg/ml chloramphenicol exposures. **F.** Plot with linear regression of 16-h cell-density values for 10 mM salicylate and for 2 mM salicylate, 8 μg/ml chloramphenicol exposures.

Since benzoate and salicylate induce multidrug resistance via the Mar regulon (24), it was of interest to test drug resistance of the evolved clones. We measured growth in chloramphenicol, an antibiotic that is effluxed by the MarA-dependent pump AcrA-AcrB-TolC (27), which confers low-level resistance. For our experiment, the same set of clones observed for growth in benzoate and salicylate were cultured in media containing 8 μg/μl chloramphenicol (**Fig. 1C, 1D**). The media were adjusted to pH 7.0 for maximal growth and contained a low concentration of benzoate or salicylate for induction of Mar regulon. Later generations (1,900–2,900 gen) reached significantly lower cell density compared to that of the ancestor W3110.

Our results suggest that populations evolving with benzoate experienced a tradeoff between benzoate-salicylate tolerance and inducible chloramphenicol resistance. This tradeoff is confirmed by the plot of benzoate tolerance versus growth in chloramphenicol (**Fig. 1E**). Clones from the ancestral strain W3110 (red circles) and early generations (orange, generation 173; yellow, generation 399) showed little growth in 20 mM benzoate, but most grew in 8 μg/ml chloramphenicol (reached OD_600_ values of at least 0.05). Middle-generation clones (light green 897, dark green 1402) grew in benzoate to higher OD_600_ and showed variable growth in chloramphenicol. By 2,900 generations (purple), all clones reached OD_600_ values of at least 0.2 in 20 mM benzoate, but the clones barely grew at all in chloramphenicol. Growth of evolved clones in salicylate, versus growth in salicylate and chloramphenicol, showed a similar reciprocal relationship (**Fig. 1F**). Outliers appeared under all conditions, as expected under selection pressure (65).

### Genome resequencing showed numerous SNPs, deletions, and IS5 insertion mutations

After 2,000 generations, 8 clones showing benzoate tolerance were chosen for genome sequencing (described under Methods). An additional 24 clones (one from each evolved population) were tested for chloramphenicol resistance; all were benzoate-tolerant compared to strain W3110. Of these 24 clones, 8 were chosen for decreased resistance to chloramphenicol. Both sets of 8 clones (16 in all) were streaked for isolation and established as strains (**Table 1**). We used the *breseq* (version 0.27.1) computational pipeline to analyze the mutation predictions of our resequenced genomes compared to the ancestral *E. coli* W3110 (lab stock D13), assembled on the W3110 reference genome (59). More than 100 mutations were detected across the sixteen sequenced genomes (**Table S1**). Mutations found in our resequenced lab stock D13 compared to the reference W3110 (**Table S2**) were filtered from the results. The types of mutations that accumulated across the sixteen strains included SNPs, small indels, insertion sequences (IS), and large IS-mediated deletions. Both coding changes and intergenic mutations were frequent. A large number of insertion knockouts were mediated by mobile insertion sequence elements such as IS5 (66). An example, *gadX*::IS5 found in clone A5-1, is shown in **Fig. 2**. The inserted sequence, including *insH* plus IS5 flanking regions, was identical to those of 11 known IS5 inserts in the standard W3110 sequence; there was also a 4-bp duplication of the target site. Insertion sequence mobility is a major source of evolutionary change in *E. coli* (67).

**FIG 2.**
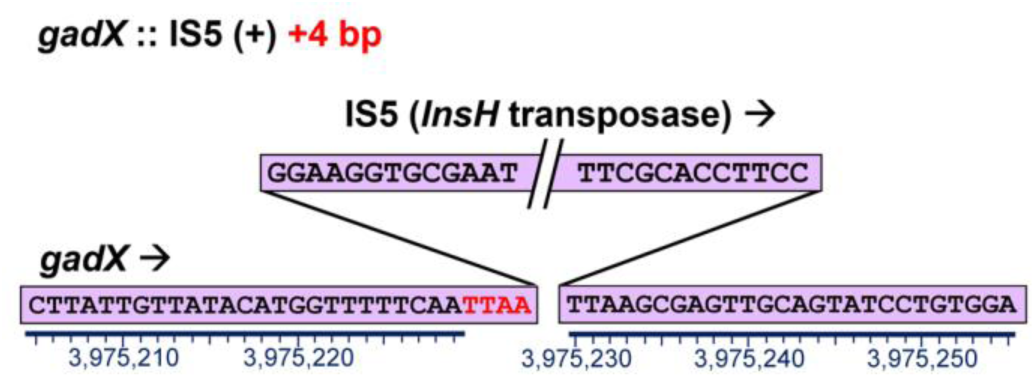
New insertion of IS5 within *gadX* including a 4-bp duplication of the target site, in the A5-1 genome at position 3,975,230.

The 2,000-generation benzoate-adapted strains were grouped in six clades based on shared mutations (**Table S1**); a representative member of each clade is shown in **Table 2**. Most of the shared mutations originated within a population, as in the case of the two strains taken from each of the A5, E1, and C3 populations; and from inadvertent cross-transfer between microplate wells. For example, one population (G5) included a strain G5-2 that shares mutations with the strains from the H1, H3 and G3 populations while sharing no mutations with strain G5-1, from the G5 population. Note also that isolated shared mutations can originate from the shared founder culture, or arise as independent genetic adaptations to a common stress condition (65).

### Mutations appeared in Mar and other multidrug efflux systems

Five of the six clades showed mutations affecting the Mar regulon, as well as other MDR genes (**Table 2**). Strain A5-1 had a 6,115-bp deletion including *marRAB* (*ydeA*, *marRAB*, *eamA*, *ydeEH*). Regulators of Mar showed point mutations in strain G5-2 (*mar* paralog *rob*, Ref. (26)) and in strain A1-1 (two-component activator *cpxA*, Ref. (68)). Additional mutations appeared in other multidrug efflux systems: *emrA*, *emrY* (32, 69), *mdtA*, deletions covering *mdtEF* (69), and *yeaS* (*leuE*) leucine export (70).

The benzoate-evolved strains were tested by MIC assay for sensitivity to the antibiotics chloramphenicol and tetracycline, whose resistance is inducible by benzoate derivatives (**Table 3**). Sensitivity was assayed in the presence or absence of a Mar inducer, 2 mM salicylate. Salicylate increased the MIC for our ancestral strain W3110, for both chloramphenicol and tetracycline. The W3110 *marA*::*kanR* construct showed less increase in MIC for chloramphenicol. For tetracycline, our *marA* knockout strain showed no loss of resistance; this may be due to induction of non-Mar salicylate-dependent resistance (24).

**Table 3.**
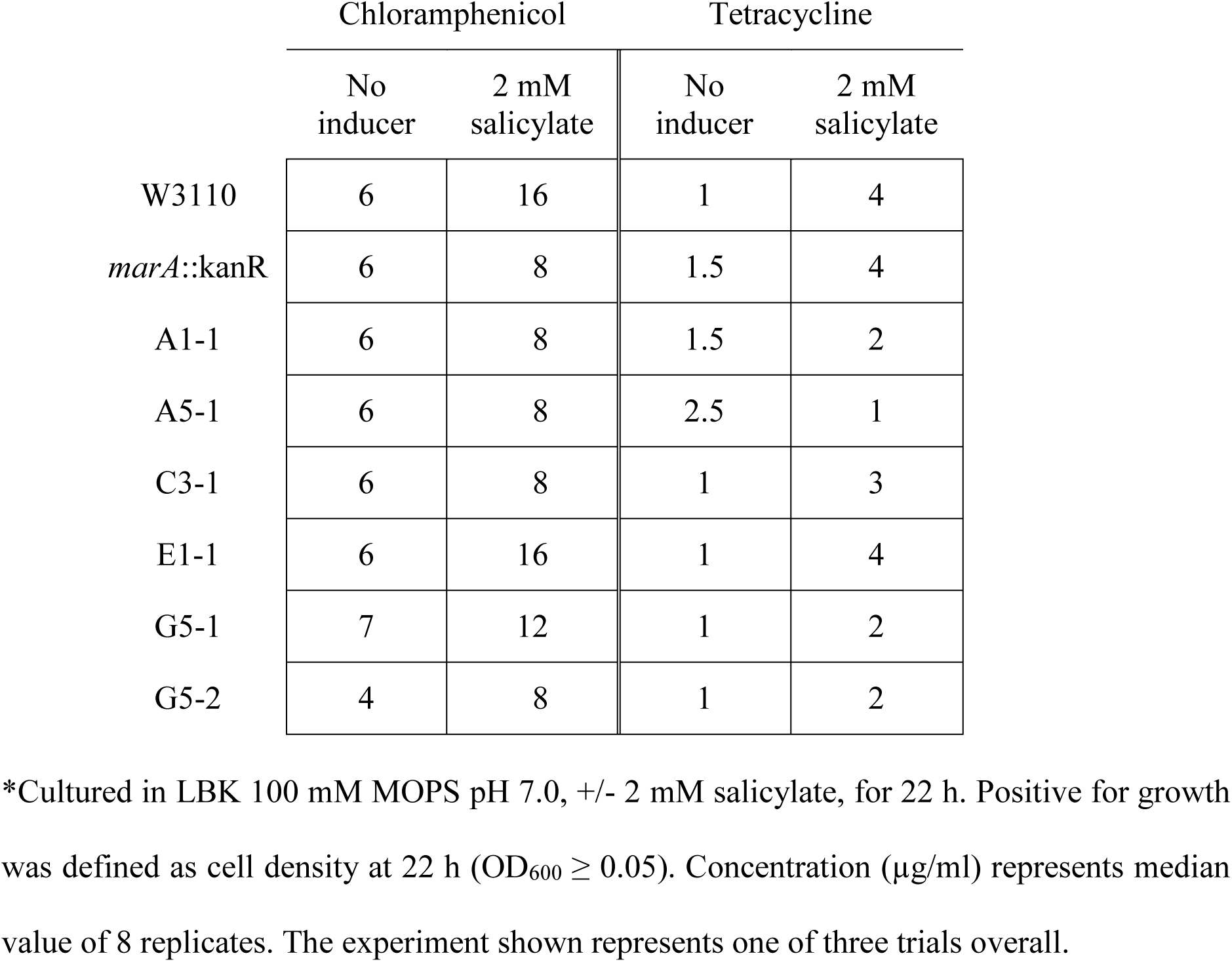
**MIC (minimum inhibitory concentration) of benzoate-evolved strains in the antibiotics chloramphenicol and tetracycline, with or without salicylate.^*^**

|  | Chloramphenicol |  | Tetracycline |  |
| --- | --- | --- | --- | --- |
|  | No inducer | 2 mM salicylate | No inducer | 2 mM salicylate |
| W3110 | 6 | 16 | 1 | 4 |
| *marA::kanR* | 6 | 8 | 1.5 | 4 |
| A1-1 | 6 | 8 | 1.5 | 2 |
| A5-1 | 6 | 8 | 2.5 | 1 |
| C3-1 | 6 | 8 | 1 | 3 |
| E1-1 | 6 | 16 | 1 | 4 |
| G5-1 | 7 | 12 | 1 | 2 |
| G5-2 | 4 | 8 | 1 | 2 |

\*Cultured in LBK 100 mM MOPS pH 7.0, +/- 2 mM salicylate, for 22 h. Positive for growth was defined as cell density at 22 h ( $OD_{600} \geq 0.05$ ). Concentration ( $\mu\text{g/ml}$ ) represents median value of 8 replicates. The experiment shown represents one of three trials overall.

In the presence of salicylate, our evolved strains A1-1, A5-1, C3-1, and G5-2 showed MIC levels half that of ancestor W3110. These lowered MIC levels were comparable to that of a *marA* mutant, and only E1-1 showed chloramphenicol resistance equivalent to that of W3110. In the absence of salicylate, the ancestral and evolved strains all showed lower MIC values due to lack of inducer. The strain G5-2 showed slightly lower MIC than that of W3110 (4 μg/m versus 6 μg/ml, respectively). Thus it is possible that G5-2 has lost an additional component of chloramphenicol resistance that does not require salicylate induction.

For tetracycline, salicylate-inducible resistance was less than that of W3110 for all of our benzoate-evolved strains with the exception of strain E1-1. The tetracycline MIC values however showed some variability; for all strains, uninducible MIC levels varied among trials from 1–3 μg/ml. For further analysis, we focused on chloramphenicol.

### Mutations appeared in Gad acid resistance, RNAP, and fimbriae

The Gad acid resistance genes are induced during cytoplasmic acidification (37) and show regulation intertwined with that of MDR systems (35). Strikingly, five of the six clades of our 16 sequenced strains showed a mutation in the Gad regulon (**Table 2**; **Table S1**). Strain E1-1 had a 14,000-bp deletion mediated by the insertion sequence *insH* flanking the *gad* acid fitness island (*gadXW*, *mdtFE*, *gadE*, *hdeDAB*, *yhiDF*, *slp*, *insH, yhiS*). Similarly, strain A1-1 showed a 10,738-bp deletion covering most of the *gad* region. A1-1 also had an insertion in the *ariR* (*ymgB*) biofilm-dependent activator of Gad (37, 71). The *mdtFE* genes encode components of the efflux pump MdtF-MdeE-TolC, which confers resistance to chloramphenicol, as well as fluoroquinolones and other drugs (72). Thus, *gad* deletion might explain chloramphenicol sensitivity of strain A1-1; but not E1-1, which was chloramphenicol resistant despite the deletion. Other strains showed mutations in the *gadX* activator: A5-1 (IS5 insertion), G5-1 (missense L199F), and G5-2 (78-bp deletion). G5-2 also showed an *hfq* point substitution, at a position known to affect function of RpoS (73), which activates Gad (2).

The *gad* mutants showed different levels of GABA production by glutamate decarboxylase (GadA) during extreme-acid exposure (incubation at pH 2) (**Fig. 3**). The two strains with full Gad deletions (A1-1 and E1-1) produced no GABA, whereas strains with *gadX* mutations (A5-1, G5-1, and G5-2) produced significantly less GABA than did the ancestor W3110 (Friedman/Conover test). Only one representative strain, C3-1, showed no Gad-related mutation; the nearest mutation was an IS5 insertion adjacent to *slp*. This strain C3-1 produced GABA in amounts comparable to those of W3110.

**FIG 3.**
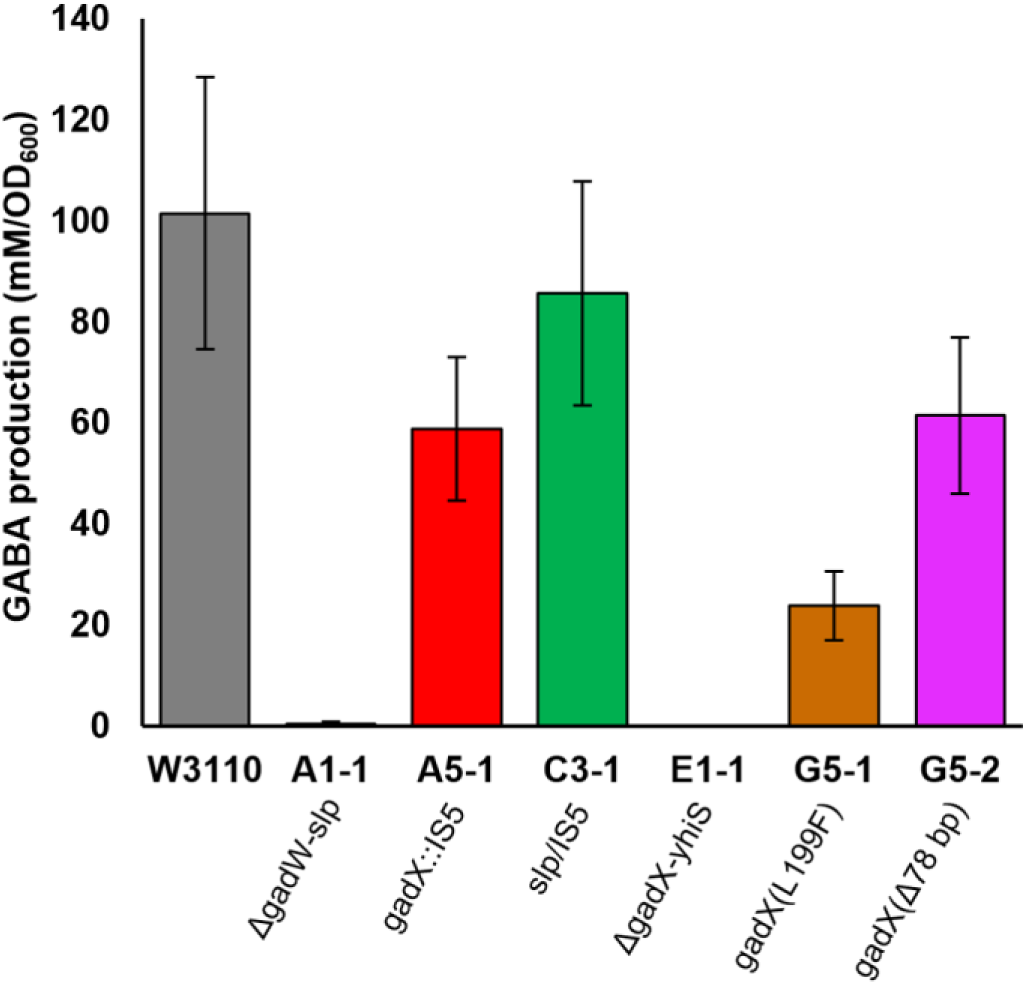
GABA produced by benzoate-evolved strains compared to W3110. Anaerobic overnight cultures in LB 10 mM glutamine, 100 mM MES pH 5.5 were adjusted with HCl to pH 2. After 2 h incubation, bacteria were pelleted and supernatant culture fluid was derivatized using EZ:faast (see Methods). GABA was quantified via GC/MS, and values were normalized to the cell density of the overnight culture. Error bars represent SEM (n=7 or 8). Genetic annotations are from Table 1.

Each benzoate-adapted strain also showed a mutation in an RNAP subunit (*rpoB, rpoA*), a sigma factor (*rpoD, rpoS*) or an RNAP-associated helicase (*hepA*). These mutations in the transcription apparatus are comparable to those we find under low-pH evolution (49). Four of the six clades had mutations in fimbria subunit *fimA* or in regulators *fimB*, *fimE*. Thus, benzoate exposure could select for loss of fimbriae synthesis. Other interesting mutations affected cell division (*ftsZ*), cell wall biosynthesis (*mrdA*), and envelope functions (*ecpD, lptD*, *ybbP, yejM*, *yfhM*, *yqiGH*, and *rfaY*). The envelope mutations suggest responses to benzoate effects on the outer membrane and periplasm.

### Benzoate-evolved strains showed increased growth rate and stationary phase cell density

In our evolution experiment, the microplate growth cycle (49) involves initially oxygenated cultures, which become semianaerobic and ultimately enter stationary phase for several hours. Thus, selection pressure occurs under changing conditions of oxygenation and cell density. Dynamic conditions are relevant to host environments such as the intestinal epithelium (74) and arterial plaque biofilms (75).

We observed the phases of growth for each benzoate-evolved strain, in order to characterize the focus of selection pressure with respect to early growth rate, stationary-phase cell density, and death phase. Observing the entire growth curve provides more information than an endpoint MIC. Growth curves were conducted in microplate wells for each of the six representative 2,000-generation benzoate-adapted strains. For each strain, eight replicate wells of the microplate were inoculated alongside eight replicate wells of strain W3110, as shown for strain G5-2 (**Fig. 4A**).

**FIG 4.**
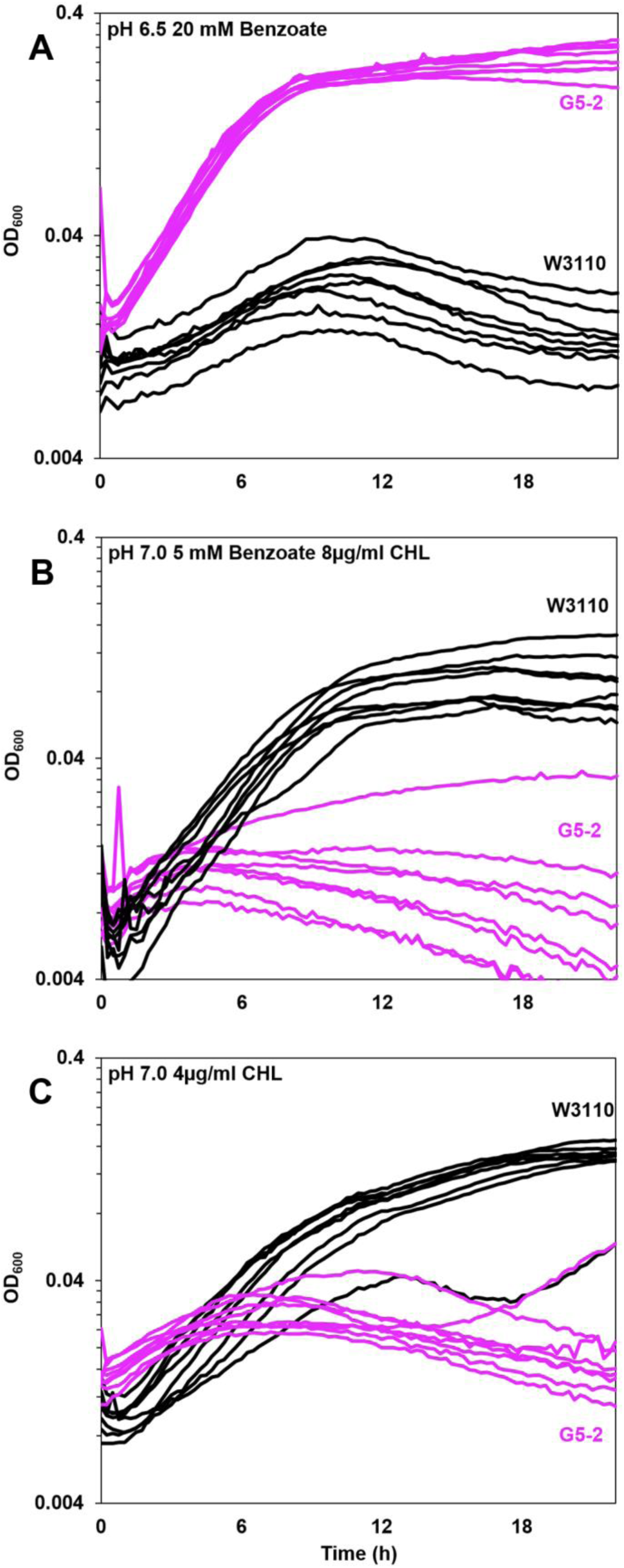
Benzoate-evolved strain G5-2 outgrows ancestor W3110 in presence of benzoate, but grows poorly in benzoate with chloramphenicol. Growth medium was LBK with (**A**) 100 mM PIPES, 20 mM benzoate pH 6.5, (**B**) 100 mM MOPS, 5 mM benzoate pH 7.0, 8 μg/ml chloramphenicol (CHL). (**C**) 100 mM MOPS, pH 7.0, 4 μg/ml CHL. For each strain, 8 replicate curves from microplate wells are shown. For strains G5-2 and W3110 in each panel, cell density values post log-phase (OD_600_ at 16 h) were ranked and compared by Friedman test; post-hoc Conover pairwise comparisons were conducted with Holm-Bonferroni adjusted p-values.

In the example shown, strain G5-2 maintained log-phase growth for approximately four hours in the presence of 20 mM benzoate (0.42±0.5 gen/h, measured over times 1–3 h). Strain G5-2 eventually reached a stationary-phase OD_600_ of approximately 1.0. By contrast, ancestral strain W3110 grew more slowly (0.18±0.01 gen/h) and peaked at OD_600_=0.5–0.7 by about 8 h. After 8 h, W3110 entered a death phase as the cell density declined. The presence of chloramphenicol, however, reversed the relative fitness of the two strains (**Fig. 4B**). The benzoate-evolved strain barely grew, and 7 of 8 replicates entered death phase by 3–4 h. By contrast, the ancestor grew steadily to an OD_600_ of 0.5–0.6, a level that was sustained for several hours.

At a lower concentration of chloramphenicol (4 μg/ml), the parental strain W3110 outgrew strain G5-2, even in the absence of benzoate inducer (**Fig. 4C**). This observation confirms the MIC result (**Table 3**) that G5-2 shows loss of an unidentified means of chloramphenicol resistance, independent of benzoate or salicylate. Other isolates showed only loss of benzoate-inducible resistance (presented below).

The effect of various permeant acids was tested, in order to determine the specificity of acid tolerance (**Fig. 5**). For all growth curves, statistical comparison was performed using cell density values at 16 h. Each panel shows a curve with median cell density (at 16 h) for a benzoate-evolved strain, as well as for strain W3110. Both benzoate and salicylate conditions showed a marked fitness advantage for all six benzoate-evolved strains (**Fig. 5A, B**). Five of the strains showed log-phase growth rates equivalent to each other, whereas G5-1 grew significantly more slowly during log phase (Friedman, Conover tests; p ≤ 0.05). All benzoate-evolved strains grew faster than strain W3110. All six benzoate-evolved strains reached equivalent plateau cell densities (OD_600_ values of approximately 1.0, with 20 mM benzoate; 0.9, with 10 mM salicylate). By contrast, in 20 mM benzoate the strain W3110 entered death phase by 10 h. This observation strongly suggests that death rates contribute to benzoate selection, besides the nominal 6.6 generations of growth per dilution cycle.

**FIG 5.**
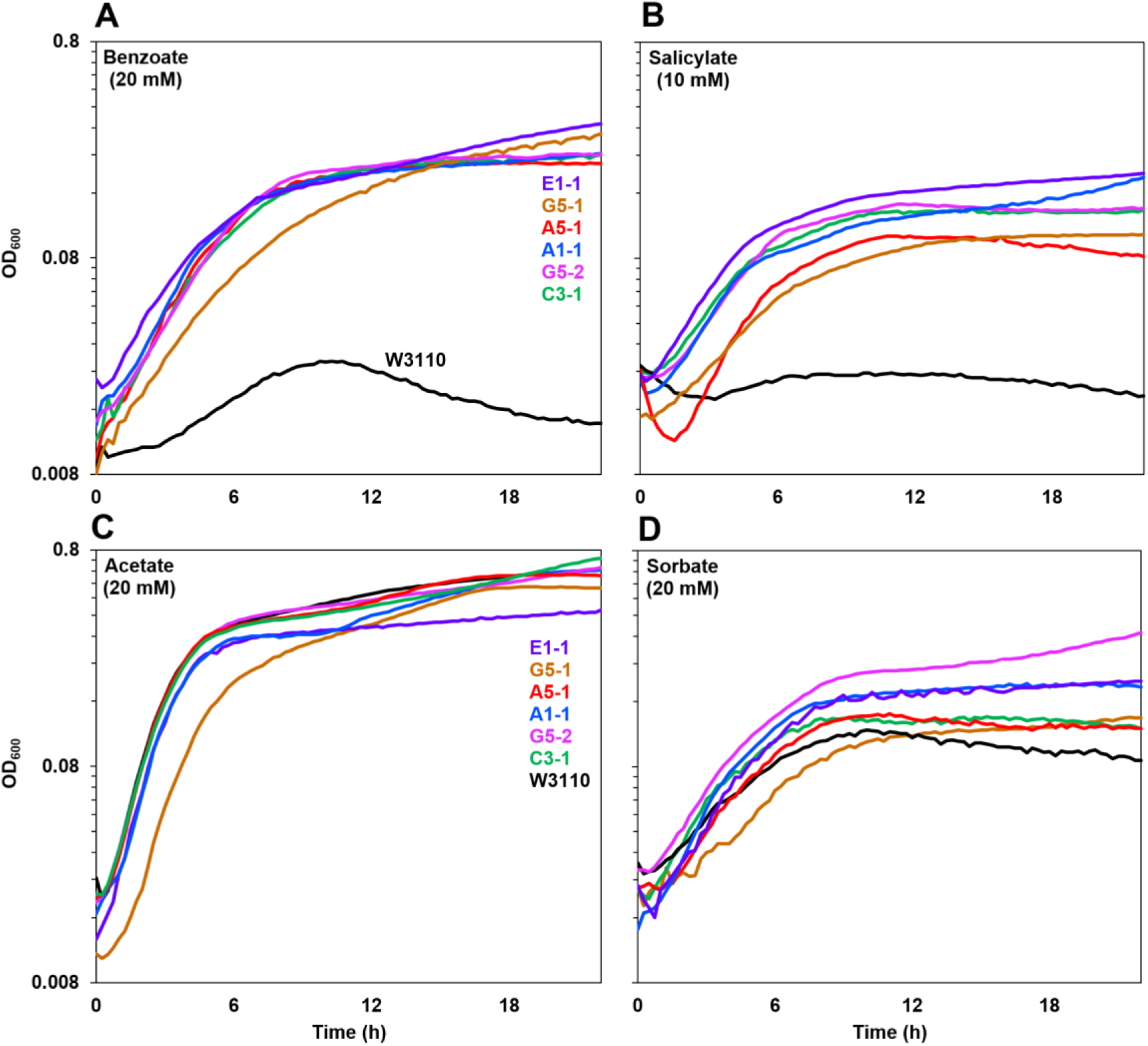
Benzoate-evolved strains all outgrow ancestor in benzoate or salicylate but not in acetate or sorbate. Growth curves of benzoate-evolved strains (colored curves) and ancestor (black curve) in LBK 100 mM PIPES pH 6.5 with (A) 20mM benzoate, (B) 10 mM salicylate, (C) 20mM acetate, or (D) 20 mM sorbate. For each strain, a curve with median cell density at 16 h is shown. Panels A and B (but not C and D) showed significantly lower 16-h cell density for the ancestral W3110 strain than for benzoate-evolved strains (Friedman test; post-hoc Conover pairwise comparisons with Holm-Bonferroni adjusted p-values).

In the presence of aliphatic acids acetate or sorbate (**Fig. 5C, D**) no significant difference was seen between growth of the ancestral strain W3110 and that of the benzoate-evolved strains. Thus, the evolved fitness advantage is unlikely to result from cytoplasmic pH depression but appears specific to the presence of aromatic acids benzoate or salicylate. Further testing with 40 μM carbonyl cyanide m-chlorophenyl hydrazone (the uncoupler CCCP) showed no difference in growth rate or stationary-phase cell density among the strains (data not shown). Thus, while decrease of proton motive force may be one factor it cannot be the sole cause of the fitness advantage of our strains.

We also tested whether the benzoate-evolved strains showed any fitness advantage with respect to pH stress. The strains were cultured in media buffered at pH7.0 (**Fig. 6A**) and at pH 4.8 or pH 9.0 (**Fig. S2**). All of the benzoate-evolved strains grew similarly to the ancestor at external pH values across the full range permitting growth. Thus, the fitness advantage of the evolved strains was specific to the presence of benzoate or salicylate.

**FIG 6.**
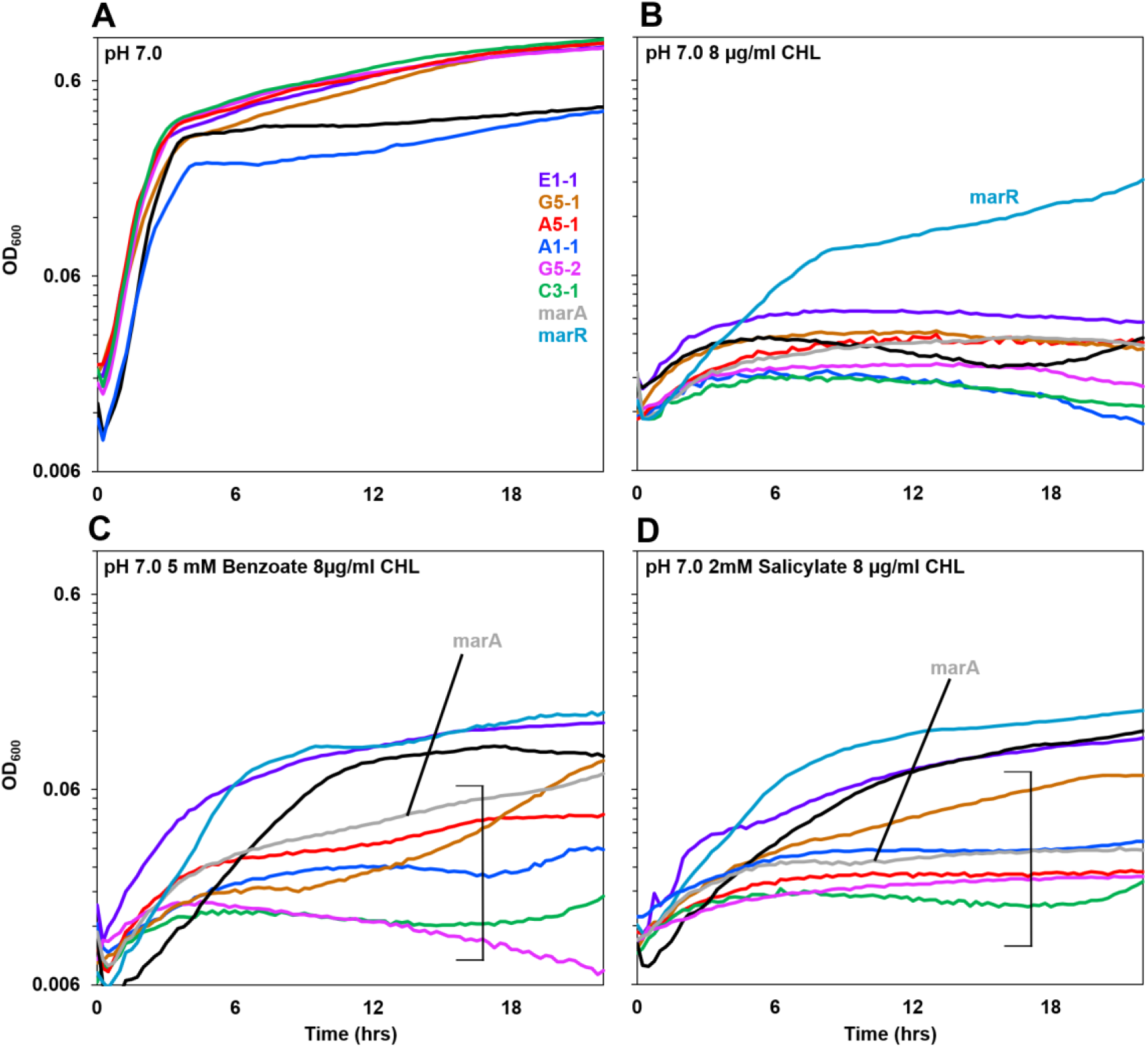
Benzoate-evolved strains are sensitive to chloramphenicol. Growth curves of benzoate-evolved strains and ancestor, compared to W3110 strains deleted for *marR* and for *marA*. Media contained LBK 100 mM MOPS pH 7.0 with (**A**) no supplements, (**B**) 8 μg/ml chloramphenicol, (**C**) 5 mM benzoate and 8 μg/ml chloramphenicol, (**D**) 2 mM salicylate and 8 μg/ml chloramphenicol. For each strain, a curve with median 16-h cell density is shown. Bracket indicates curves with cell density at16 hours that was lower than the cell density of ancestral strain W3110 (Friedman test; post-hoc Conover pairwise comparisons with Holm-Bonferroni adjusted p-values). For panels C and D, **Figure S3** shows all replicates of each strain plotted individually.

### Chloramphenicol inhibits growth of benzoate-evolved clones, despite benzoate fitness advantage

Since each of the six clades showed a mutation in an MDR gene or regulator (discussed above), we characterized the growth profiles of all strains in the presence of chloramphenicol (**Fig. 6B, C, D**). In the absence of benzoate or salicylate inducer (**Fig. 6B**), all strains showed growth curves equivalent to that of W3110 or W3110 *marA*::*kanR* (which lacks the MarA activator of MDR efflux). Only the *marR*::*kanR* strains (constitutive for activator *marA*) showed resistance. In the presence of benzoate (**Fig. 6C**) or salicylate (**Fig. 6D**), the various benzoate-evolved strains reached different cell densities in the presence of chloramphenicol. Panels presenting all 8 replicates of each strain are presented in supplemental **Figures S3** and **S4**.

In the presence of chloramphenicol (with benzoate or salicylate) only strain E1-1 consistently grew at a rate comparable to W3110 (**Fig. 6C, D**). Strains A5-1 and G5-1 grew to a lower density, comparable to that of the *marA*::*kanR* strain. Strains C3-1 and G5-2 showed hypersensitivity to chloramphenicol, with cell densities significantly below that of *marA*::*kanR*. The sensitivity of strain C3-1 is noteworthy given the absence of Mar or Gad mutations. Another strain, G5-2, shows chloramphenicol sensitivity greater than the level that would be predicted from loss of Rob activating MarA (25). Thus, the C3-1 and G5-2 genomes may reveal defects in other benzoate-inducible MDR genes, as yet unidentified.

### Mutations in *rob*, *gadX*, and *cpxA* do not affect chloramphenicol sensitivity or benzoate tolerance

Since five of the six benzoate-evolved strains showed defects affecting Gad regulon, we tested the role of *gadX* in chloramphenicol resistance (**Fig. 7**). In the W3110 background, a *gadX*::*kanR* knockout (green) showed more sensitivity to chloramphenicol, as did the construct W3110 *marA*::*kanR* (gray). Thus it is possible that GadX has some uncharacterized role in chloramphenicol efflux, either via activation of the Mar regulon, or else via activation of *mdtE*, *mdtF* in the Gad fitness island (36).

**FIG 7.**
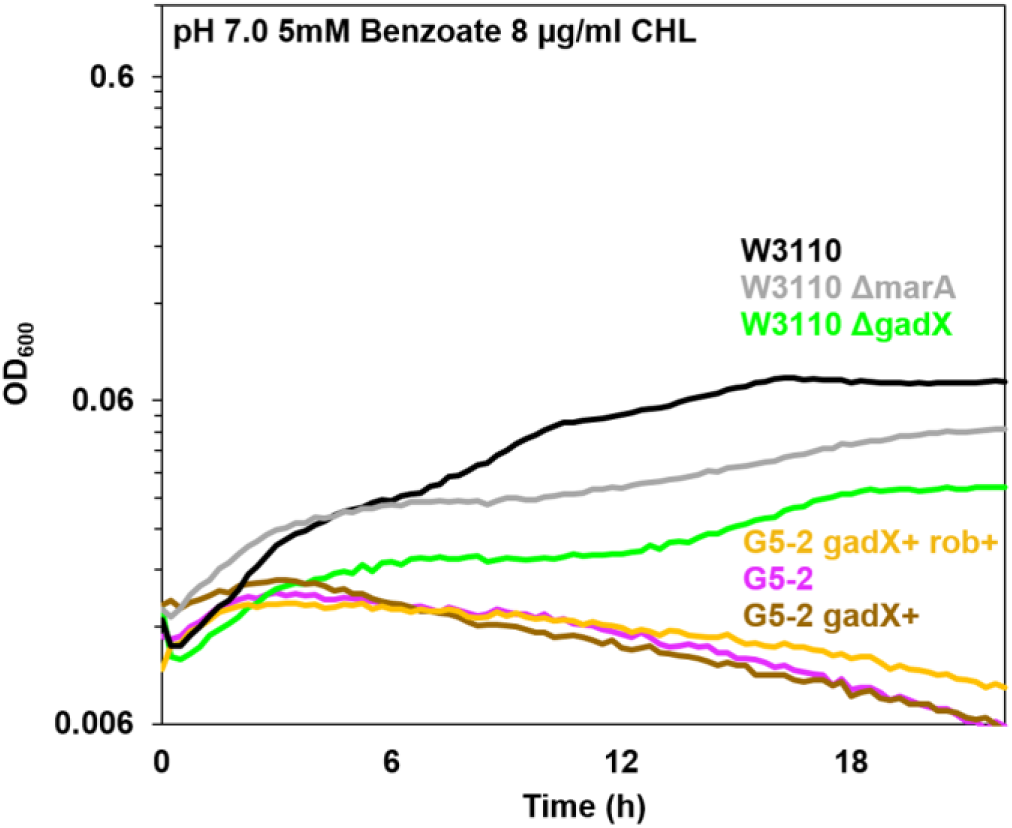
W3110 *gadX*::*kanR* shows sensitivity to chloramphenicol. Growth curves were conducted in LBK 100 mM PIPES pH 7.0 with 5 mM benzoate, 8μg/ml chloramphenicol, as for Figure 6. At 16 h, the *gadX*::*kanR* knockout and the *marA*::*kanR* strain both grew significantly less than strain W3110. The cell density of strain G5-2 showed no difference from those of G5-2 *gadX*+ or of G5-2 *gadX*+ *rob*+ (Friedman test; post-hoc Conover pairwise comparisons with Holm-Bonferroni adjusted p-values).

We also tested the role of *gadX* and *rob* in chloramphenicol sensitivity of the benzoate-evolved strain G5-2. The G5-2 mutant alleles of Mar activator *rob* (S34P) and Gad activator *gadX* (Δ78bp) were replaced by cotransduction. The ancestral alleles of *rob* and of *gadX* were each moved into G5-2 by cotransduction with linked markers *yjjX790*::*kanR* and *treF774*::*kanR* respectively. Constructs of G5-2 with allele replacements *gadX^+^* and *rob*^+^ *gadX*^+^ were cultured with chloramphenicol in the presence of 5 mM benzoate (**Fig. 7**). Both constructs were as sensitive to chloramphenicol as the parental G5-2 strain. Furthermore, strain G5-2 constructs with either *rob*^*+*^, *gadX*^*+*^, or *rob*^+^ *gadX*^+^ showed no significant loss of benzoate tolerance compared to the parental strain G5-2 (data not shown). Thus, both the benzoate fitness advantage and the chloramphenicol sensitivity of G5-2 must involve other unidentified mutations in addition to *rob* or *gadX*.

We also tested the effect of *cpxA*^*+*^ on the benzoate fitness and chloramphenicol sensitivity of strain A1-1. The parental allele of MDR regulator *cpxA* was moved from W3110 into A1-1, replacing *cpxA*(N107S) with a marker *fdhD*::*kanR* linked to *cpxA*^*+*^ (**Fig. S5**). Growth curves were conducted as for **Figures 5** **and** **6**, tested under the conditions of 20 mM benzoate pH 6.5; benzoate with chloramphenicol; and salicylate with chloramphenicol. Under all conditions, the cell density of strain W3110 *cpxA*::*kanR* (orange) showed no difference from that of W3110; and strain JLS1607 (A1-1 *fdhD75*8::*kanR cpxA+*) (brown) showed no difference from JLSK0001 (A1-1) (blue). Thus, strain A1-1 must also contain benzoate-selected mutations in other unknown MDR genes.

## DISCUSSION

In order to identify candidate genes for benzoate stress response, we sequenced the genomes of experimentally evolved strains (41, 42, 46, 49, 76). We observed over 100 distinct mutations in the sequenced isolates after 2,000 generations, including a surprising number of knockout alleles due to mobile elements (66, 67) in particular IS5 insertions and IS5-mediated deletions (52). The *E. coli* K-12 W3110 genome contains many IS-elements, including eight copies of IS1, five copies of IS2, and copies of other less well-studied IS types where the most prevalent are IS1 and IS5 (59, 77, 78). Transposition of IS5 may be induced by environmental factors such as motility conditions, which induce IS5 insertion upstream of motility regulator *flhD* (79). Nonetheless, finding an additional 25 insertion sequences under benzoate selection is remarkable. Our detection of such IS inserts was enabled by use of the updated *breseq* pipeline (53).

Benzoate exposure decreases the cell’s PMF while simultaneously upregulating several regulons, including many involved in drug resistance. Our data suggests that benzoate exposure selects for genetic changes in *E. coli* that result in, over time, the loss of energetically costly systems such as Mar and other MDR regulons, as well as Gad acid-inducible extreme-acid regulon. MarA is a potent transcriptional factor in *E. coli*, upregulating numerous efflux pumps and virulence factors (8, 25, 27). The transcription and translation of so many gene products would result in a considerable energy strain on the individual cell.

Many MDR complexes are efflux pumps that spend PMF, which is diminished by partial uncouplers such as benzoate or salicylate. The decreased energy expense could Mar gene products, such as those we see in A5-1 and G5-2. At least one strain (G5-2) shows detectable loss of chloramphenicol resistance in the absence of Mar inducer, suggesting loss of constitutive MDR as well as the inducible system. Thus it might be possible for benzoate and salicylate to select against a broad spectrum of drug resistance systems.

A similar energy load may occur under benzoate-depressed cytoplasmic pH, where the Gad regulon is induced. Gad includes expression of numerous gene products such as glutamate decarboxylase, whose activity enhances fitness only in extreme acid (pH 2) (35) and which breaks down valuable amino acids. Thus the deletion of the *gad* region (seen in strains A1-1 and E1-1) could eliminate a fruitless energy drain. For comparison, we find similar Gad deletions in our pH 4.6 evolution experiment (He et al. manuscript in preparation) but not in our evolution experiment conducted at pH 9.2 (unpublished). This implies that both pH 4.6 and benzoate/pH 6.5 induce Gad under conditions where glutamate decarboxylase fails to help the cell, and thus the energy-expensive Gad expression is selected against.

The Gad region also includes genes encoding an MDR complex (*mdtEF*) (36); and a *gadX* knockout showed some loss of chloramphenicol resistance (**Fig. 7**). Note however that our E1-1 strain retains chloramphenicol resistance despite Gad deletion, whereas strain C3-1 (chloramphenicol sensitive) possesses the entire Gad region except for a possible defect in *slp*, encoding an acid-resistance outer membrane protein. Thus, Gad mutations alone cannot explain the chloramphenicol sensitivities of our strains.

Another possible consequence of energy stress is the loss of fimbriae synthesis (strains A5-1, C3-1, E1-1, G5-1). Avoiding fimbriae production could save energy for benzoate-stressed cells. Fim genes also show deletion under evolution at pH 9.2 (Issam Hamdallah, unpublished), but not at pH 4.6. This suggests a hypothesis that fimbriae biosynthesis is dependent less on pH than on the protonmotive force (PMF), which is depleted both by benzoate and at high external pH, a condition that decreases PMF (5).

The progressive loss of antibiotic resistance is a remarkable consequence of benzoate selection, evident as early as 1,500 generations (**Fig. 1**). Several observations point to the existence of inducible MDR systems yet to be discovered. Strains C3-1 and G5-2 show hypersensitivity to chloramphenicol, beyond the level of sensitivity seen in a *marA* knockout (**Fig. 6C, D**; **Fig. S3**). Furthermore, the reversion of mutant alleles of *rob*, *gadX*, and *cpxA* do not diminish the phenotypes of the 2,000-generation strains. It is likely that these alleles conferred a fitness advantage early on in our evolution experiment (65) but have since been superseded by further mutations in as yet unidentified players in drug resistance.

The fitness tradeoff between drug resistance and benzoate/salicylate exposure has implications for the human microbiome. Mar and homologs such as Mex are reported in numerous bacteria, including proteobacteria and *Bacteroides fragilis* (80, 81). Salicylate is a plant defense molecule commonly obtained via human diets rich in fruits and vegetables (82, 83). Aspirin is deacetylated in the liver and stomach, forming salicylic acid, the primary therapeutic agent (84). As a membrane-permeant acid, salicylic acid permeates human tissues nonspecifically. Both food-related and aspirin-derived salicylates come in contact with enteric gut bacteria, where they would be expected to activate Mar-like antibiotic resistance systems. Commonly prescribed for cardiac health, aspirin releases salicylate at plasma levels of approximately 0.2 mM (16, 19, 20, 22).

Intestinal salicylate levels are poorly understood, but even lower concentrations of this antimicrobial agent could have fitness effects. For comparison, small concentrations of antibiotics, well below the MIC, can select for resistance (85, 86). Similarly, it may be that small concentrations of a resistance-reversing agent such as salicylate have a significant fitness cost for MDR bacteria. Furthermore, the effective concentration of permeant acids such as salicylate is amplified exponentially by the pH difference across the bacterial plasma membrane. Even mild acidity in the intestinal lumen (pH 6–6.5) could amplify the bacterial cytoplasmic concentration of a permeant acid by 10- to 30-fold.

Aspirin therapy is known to prevent clotting by inactivation of human cyclooxygenase, leading to suppression of prostaglandins. There is little attention, however, to the possible effects of aspirin on human-associated bacteria. Gram-negative pathogens such as *Pseudomonas* are found in arterial plaques and associated with heart attacks (75). Aspirin-derived salicylate in plasma might provide a fitness cost for such bacteria. Aspirin also prevents colon cancer, by some unknown mechanism (16, 20). Colon cancer depends on colonic bacteria and the formation of biofilms (87, 88).

Long-term salicylate exposure via aspirin therapy may select a microbiome that is salicylate-tolerant but drug-sensitive. A salicylate-adapted microbiome may confer the benefit of excluding drug-resistant pathogens that lack salicylate tolerance. In blood plasma, salicylate levels might help exclude bacteria from arterial plaques. An adverse consideration, however, is that the salicylate-adapted microbiome of the colon may be more vulnerable to high dose antibiotic therapy. For the future, we are testing these speculative possibilities in host microbial models.

## Acknowledgements

We thank Zachary Blount, Jeff Barrick, Michael Harden, Rohan Maddamsetti, Peter Lund, and Bradley Hartlaub for valuable discussions. We thank Anna Tancredi for testing the *gadX* mutant. This work was funded by grant MCB-1329815 from the National Science Foundation, and by the Kenyon College Summer Science Scholars.

